# Testicular Dysgenesis Syndrome-like morphology and gene expression, and activation of Hypoxia Inducible Factor 1 Alpha in juvenile lamb testes following developmental exposure to low-level environmental chemical mixture

**DOI:** 10.1101/2022.02.08.479595

**Authors:** Chris S. Elcombe, Ana Monteiro, Matthew R. Elcombe, Mohammad Ghasemzadeh-Hasankolaei, Neil P. Evans, Michelle Bellingham

**Author notes:** Post-publication corresponding author: Michelle Bellingham Room 242, Jarrett Building, Garscube Estate, University of Glasgow, G61 1QH. Pre-publication corresponding author: Chris Elcombe Room 234D, Jarrett Building, Garscube Estate, University of Glasgow, G61 1QH.

## Abstract

Current declines in male reproductive health may, in part, be driven by anthropogenic environmental chemical exposure. Using the biosolid treated pasture (BTP) sheep model, this study examined the effects of gestational exposure to a translationally relevant mixture of environmental chemicals. At 8 weeks of age, ram lambs gestationally exposed to BTP were lighter than control, and their testes contained fewer germ cells and had a greater proportion of Sertoli-cell-only seminiferous tubules. Effects of biosolid exposure on the testicular transcriptome correlated with human testicular dysgenesis syndrome (TDS) patient data. Common differentially expressed genes indicated changes in apoptotic and mTOR signalling, which concurs with previous gene expression data from neonatal BTP lambs. Gene expression data and immunohistochemistry indicates increased HIF1α activation and nuclear localisation in BTP exposed animals, known to disrupt testosterone synthesis. Together, these results provide a potential mechanism for the pathogenesis of this phenotype, and TDS in humans.

## 1. Introduction

Male reproductive health has been reported to be in decline for the past 80 years ^1^, with reduced semen quality and fecundity, and increased incidence of testicular dysgenesis syndrome (TDS; encompassing cryptorchidism, hypospadias, hypogonadism, infertility, and testicular germ cell cancer)^2^. While precise mechanisms behind the pathogenesis of TDS are still to be fully elucidated, a link has been made to foetal Sertoli and Leydig cell dysfunction ^3,4^. While many contributory factors to the decline in male fertility have been identified, including improper nutrition, sedentary lifestyle, and stress, much attention has focused on the role of environmental chemicals (ECs), in particular endocrine disrupting chemicals (EDCs) ^5–7^. A variety of ECs and families of ECs are known to be able to adversely affect testicular development, for example, gestational exposure to phthalates can negatively affect germ cell development, sperm motility, and testosterone production ^8,9^. In addition to such studies which have examined the effects of individual ECs, several studies have used component-based methodologies to examine the effects of low dose mixtures of ECs. When presented in this form, it is noteworthy that adverse effects of EC exposure have been reported even when individual chemicals were present at doses at or below their respective tolerable daily intake (TDI) values ^10,11^. This exposure scenario is of note as it more realistically reflects human EC exposure, which is characterised as chronic, extremely complex, and very low-level.

The human EC exposome is represented in our wastewater. The solids from wastewater treatment, biosolids, are extensively used as an agricultural fertiliser and reflect the human EC exposome in terms of complexity and concentration ^12–16^. ECs can be measured in tissue samples collected from sheep pastured on biosolid treated pasture (BTP) ^12,17–20^, as well as the organs of their offspring ^12,19,20^. These ECs include alkylated phenols, dioxin-like compounds, flame retardants such as polychlorinated biphenyls (PCBs) and polybrominated diphenyl ethers (PBDEs), pharmaceuticals and personal care products (PPCPs), plasticising agents such as phthalates and bisphenol A (BPA), polycyclic aromatic hydrocarbons (PAHs), and metabolites thereof ^13–15^. Experiments using the BTP sheep model have previously shown that *in utero* EC exposure can cause a multitude of effects in offspring. These include altered behaviour, differences in bone composition, disruption to cellular and hormonal processes, and changes in liver function, as well as effects on gonadal development in males and females ^17,18,21–30^. With specific regards to male gonadal development, gestational exposure to BTPs has been associated with reduced testicular weights and fewer gonocytes, Leydig cells, and Sertoli cells in 110-day-old foetuses ^23^. Neonatal (1-day-old) lambs also display fewer gonocytes as well as an increased incidence of Sertoli-cell-only (SCO) seminiferous tubules ^24^. This study indicated that there may be two phenotypic responses to EC exposure within the male lambs, one more susceptible to disruption and another resistant. This observation mirrors findings in 9-month-old offspring where a subset of animals was identified that had reduced germ cell numbers and an increased incidence of SCO seminiferous tubules ^17^. Puberty in male sheep begins at approximately 8 weeks of age, therefore in this study the morphology and transcriptome of gestationally BTP exposed juvenile (8-week-old) ram lamb testes were examined to provide insights into the mechanisms underlying observed adverse morphological and functional outcomes.

## 2. Results

### 2.1. Anatomical and histopathological analyses

Mean morphological indices are presented in Table 1. Mean body weight was significantly (p = 0.010) lower in biosolid compared to control lambs. Mean pituitary weight was significantly (p = 0.042) lower in biosolid (0.435 ± 0.054 g) compared to control (0.489 ± 0.060 g) lambs, but when corrected for body weight there was a trend (p = 0.054) for pituitary/BWT to be higher in biosolid lambs. Mean adrenal weight was not affected by treatment but when expressed relative to body weight was significantly (p = 0.038) higher in biosolid compared to control lambs.

**Table 1.** Anatomical metrics for control and biosolid exposed juvenile ram lambs. Data is presented as mean ± SD for animal body weights, absolute organ weights, and relative organ to body weights. All p-values derived from generalised linear models of biosolid animals compared with controls. Bold italics indicates p<0.05.

| Morphological indices | Control (n=11) |  |  | Biosolid (n=11) |  |  | p-value |
| --- | --- | --- | --- | --- | --- | --- | --- |
| ***Body Weight (BWT) (kg)*** | ***25.227*** | ***±*** | ***3.629*** | ***21.000*** | ***±*** | ***3.369*** | ***0.010*** |
| Gonads (g) | 43.993 | ± | 12.538 | 35.270 | ± | 12.296 | 0.115 |
| Testes (g) | 27.690 | ± | 8.926 | 21.191 | ± | 7.700 | 0.089 |
| Epididymides (g) | 16.269 | ± | 4.899 | 14.079 | ± | 5.197 | 0.334 |
| Adrenals (g) | 1.914 | ± | 0.374 | 2.148 | ± | 0.457 | 0.203 |
| Thyroids (g) | 2.038 | ± | 0.648 | 1.748 | ± | 0.439 | 0.233 |
| ***Pituitary (g)*** | ***0.489*** | ***±*** | ***0.060*** | ***0.435*** | ***±*** | ***0.054*** | ***0.042*** |
| Gonads / BWT (g/kg) | 1.756 | ± | 0.459 | 1.680 | ± | 0.598 | 0.743 |
| Testes / BWT (g/kg) | 1.111 | ± | 0.327 | 1.003 | ± | 0.340 | 0.469 |
| Epididymides / BWT (g/kg) | 0.653 | ± | 0.180 | 0.677 | ± | 0.277 | 0.818 |
| ***Adrenals / BWT (g/kg)*** | ***0.079*** | ***±*** | ***0.026*** | ***0.105*** | ***±*** | ***0.029*** | ***0.038*** |
| Thyroids / BWT (g/kg) | 0.082 | ± | 0.025 | 0.085 | ± | 0.026 | 0.751 |
| ***Pituitary / BWT (g/kg)*** | ***0.019*** | ***±*** | ***0.002*** | ***0.021*** | ***±*** | ***0.019*** | ***0.054*** |

Representative immunohistochemical images from control and biosolid testes are shown in Figure 1(A and B). The mean relative germ cell: Sertoli cell ratio (Figure 1C) was significantly (p = 0.009) lower in biosolid lambs (0.67 ± 0.22) compared to controls (1 ± 0.29). The mean percentage of SCO seminiferous tubules (Figure 1D) was significantly (p = 0. 007) higher in biosolid lambs (12.56 ± 11.49%) compared to control (2.27 ± 1.81 %). The mean minimum Feret’s diameter of seminiferous tubules (Figure 1E) was significantly (p = 0.021) smaller in biosolid (77.51 ± 16.51 μm) than in control (94.89 ± 13.36 μm) lambs.

**Figure 1.**
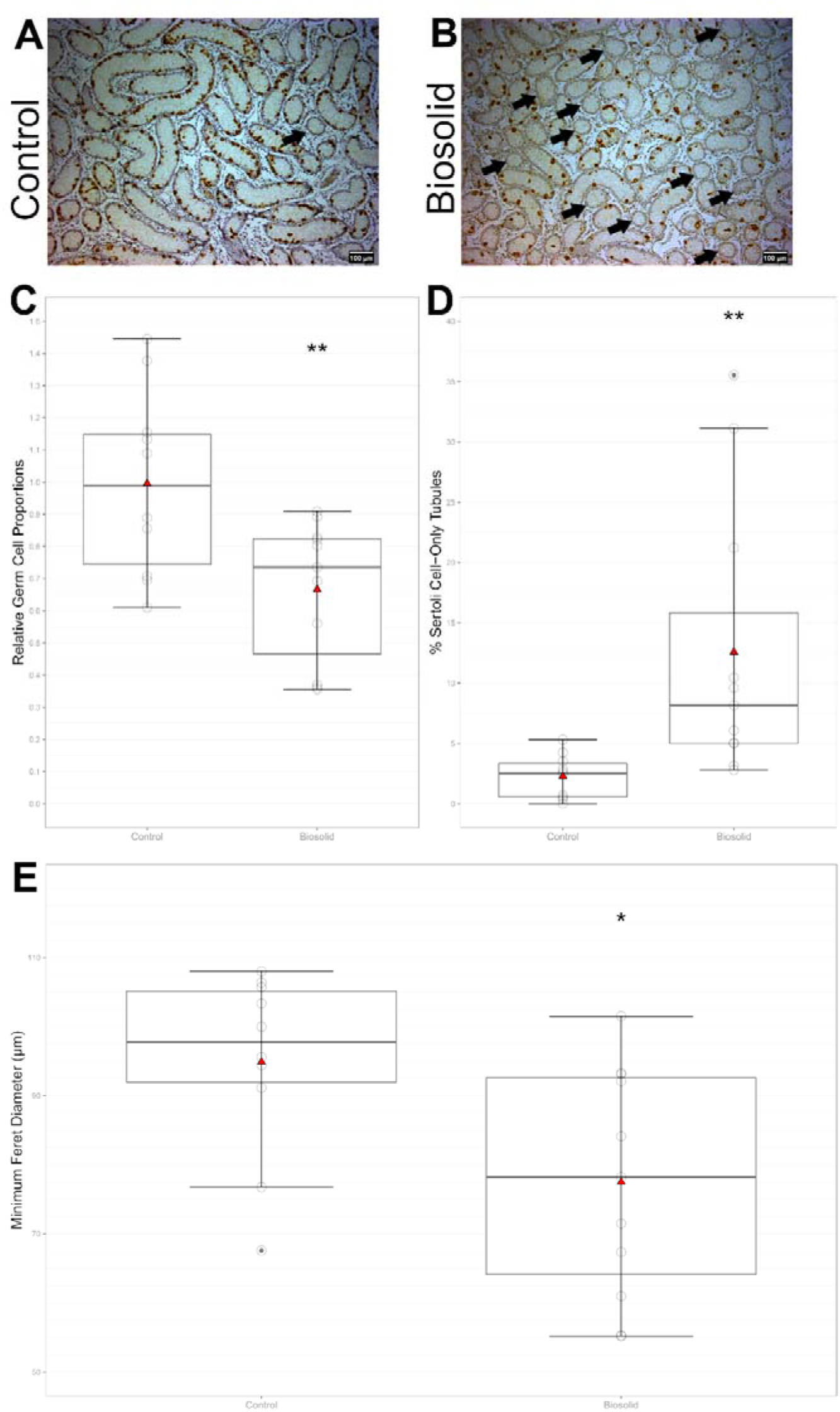
Histopathological findings in juvenile control and biosolid lamb testes. Representative images of H&E and DDX4-DAB stained tissue sections from 8-week-old control (A) and biosolid (B) lambs, taken at 40x magnification. Arrows indicate Sertoli-cell-only (SCO) seminiferous tubules. Relative proportions of germ cells to Sertoli cells (C) and percent of tubules which were SCO (D) (p = 0.009 and p = 0.007 respectively). Seminiferous tubule minimum Feret’s diameters (p=0.021) (E). Boxes represent 25^th^ to 75^th^ percentile, horizontal bar indicates 50th percentile, whiskers indicate range excluding outliers, solid filled circles show outliers, open circles show individual data points, and red triangles show means.

### 2.2. Gene expression and gene ontology analyses

DGE analysis identified 1382 DEGs between the biosolid and control groups (726 with higher expression, 656 with lower). 99 DEGs had an FDR ≤ 0.1 (60 with higher expression, 39 with lower). A z-score heatmap of these genes is shown in Figure 2 (Differential expression data presented in Supplementary Data 2). GO analysis of the 99 DEGs indicated 6 GO terms as significantly (p < 0.05) enriched (Supplementary Data 3). Within these identified GO terms, expression levels of 2 genes showed significant (p < 0.05) correlation with the geometric mean of germ cell markers, both from different GO term groups (Supplementary Data 4). These represent 12.5% and 50% of genes within respective GO terms (Nucleic acid binding and Protein ADP-ribosylation, respectively), and 2.94% of total genes identified.

**Figure 2.**
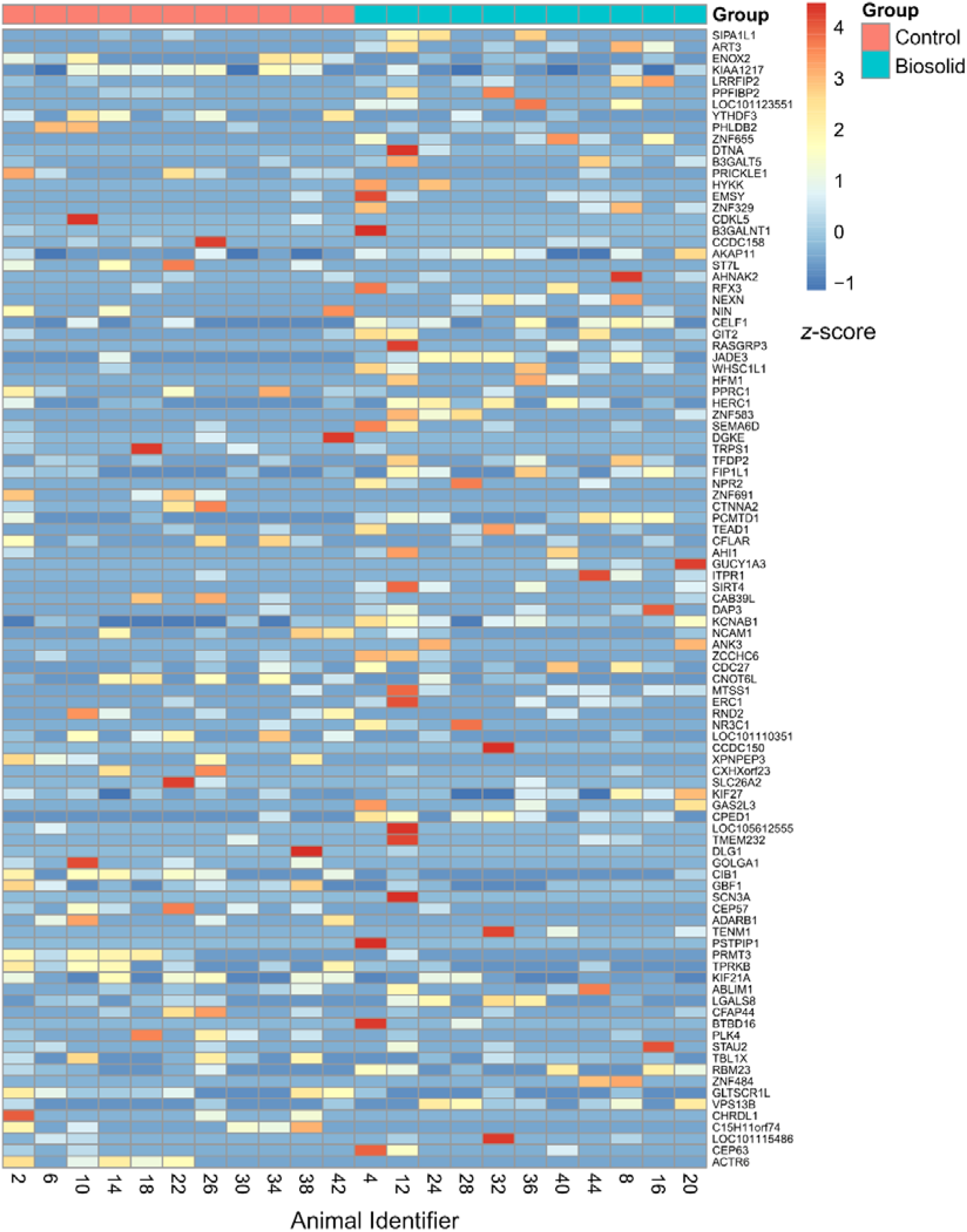
Heat map of DEGs plotted against z-score. Genes are ordered from top to bottom by increasing p-value.

### 2.3. BaseSpace correlations

A significant (p < 0.0001) positive correlation was identified between the biosolid DEG data and 7 published DEG data sets from 3 separate human testes gene expression studies (Supplementary Data 5). A gene list was created by identification of DEGs in common across each dataset per study and then combining between studies. This list comprised of 520 genes and was submitted to DAVID for GO and pathway analysis. 9 KEGG pathways and 25 GO terms were identified as significantly (p < 0.05) enriched (Table 2). Within these pathways and terms, 6 genes showed significant (p < 0.05) correlation with the geometric mean of germ cell markers, each from different GO term groups **(Error! Reference source not found.** Supplementary Data 6). This represented between 9% and 33% of genes within respective GO terms, and 1.3% of total genes identified.

**Table 2.** GO terms and KEGG pathways identified as potentially enriched by DAVID within the list of common DEG between the biosolid data and at least one correlating study identified by BaseSpace.

| Category | Term | Fold Enrichment | p-value |
| --- | --- | --- | --- |
| GO: Cellular Component | Cytosol | 2.00 | 0.000030 |
| GO: Molecular Function | Zinc ion binding | 1.75 | 0.000140 |
| GO: Cellular Component | Nucleolus | 2.04 | 0.000285 |
| GO: Cellular Component | Intracellular ribonucleoprotein complex | 6.03 | 0.000926 |
| GO: Molecular Function | mRNA 3'-UTR binding | 7.51 | 0.000999 |
| KEGG | Cell cycle | 3.52 | 0.001055 |
| GO: Cellular Component | Early endosome | 3.16 | 0.002503 |
| GO: Molecular Function | Hydrolase activity | 3.98 | 0.003734 |
| KEGG | Purine metabolism | 2.76 | 0.004030 |
| GO: Biological Process | Apoptotic process | 3.88 | 0.004223 |
| KEGG | Metabolic pathways | 1.45 | 0.007357 |
| GO: Biological Process | mTOR signalling | 8.96 | 0.008819 |
| KEGG | Lysine degradation | 4.51 | 0.010057 |
| GO: Molecular Function | GTPase activator activity | 2.61 | 0.014330 |
| KEGG | ErbB signalling pathway | 3.33 | 0.018098 |
| GO: Biological Process | Ras protein signal transduction | 4.70 | 0.020505 |
| GO: Molecular Function | DNA-directed DNA polymerase activity | 6.59 | 0.021242 |
| GO: Biological Process | Positive regulation of gene silencing by miRNA | 12.48 | 0.021909 |
| GO: Cellular Component | mTORC2 complex | 11.95 | 0.024152 |
| GO: Molecular Function | DNA-dependent ATPase activity | 5.96 | 0.027828 |
| GO: Biological Process | Mitochondrial fragmentation involved in apoptotic process | 10.92 | 0.028559 |
| GO: Biological Process | Protein localization to microtubule | 10.92 | 0.028559 |
| KEGG | Pyrimidine metabolism | 2.98 | 0.029146 |
| GO: Biological Process | Platelet-derived growth factor receptor signalling pathway | 5.82 | 0.029464 |
| GO: Molecular Function | RNA polymerase II core promoter proximal region sequence-specific DNA binding | 1.92 | 0.030620 |
| GO: Cellular Component | Nucleus | 1.25 | 0.032457 |
| GO: Biological Process | Retrograde transport, endosome to Golgi | 4.04 | 0.033621 |
| KEGG | Phosphatidylinositol signalling system | 2.86 | 0.034587 |
| GO: Biological Process | Peptidyl-tyrosine autophosphorylation | 9.71 | 0.035899 |
| GO: Molecular Function | ATP binding | 1.32 | 0.036382 |
| KEGG | Biosynthesis of antibiotics | 2.07 | 0.039467 |
| GO: Biological Process | Negative regulation of viral transcription | 8.74 | 0.043875 |
| KEGG | Base excision repair | 5.01 | 0.044115 |
| GO: Biological Process | Mitochondrion organization | 4.85 | 0.047341 |

### 2.4. Quantitative PCR

Of the 13 genes quantified by qPCR, 4 had significantly (p < 0.05) higher expression in biosolid animals than in control (Figure 3): *FASN* (p = 0.045), *HK1* (p= 0.026), *PDPK* (p = 0.042), and *VEGFA* (p = 0.037).

**Figure 3.**
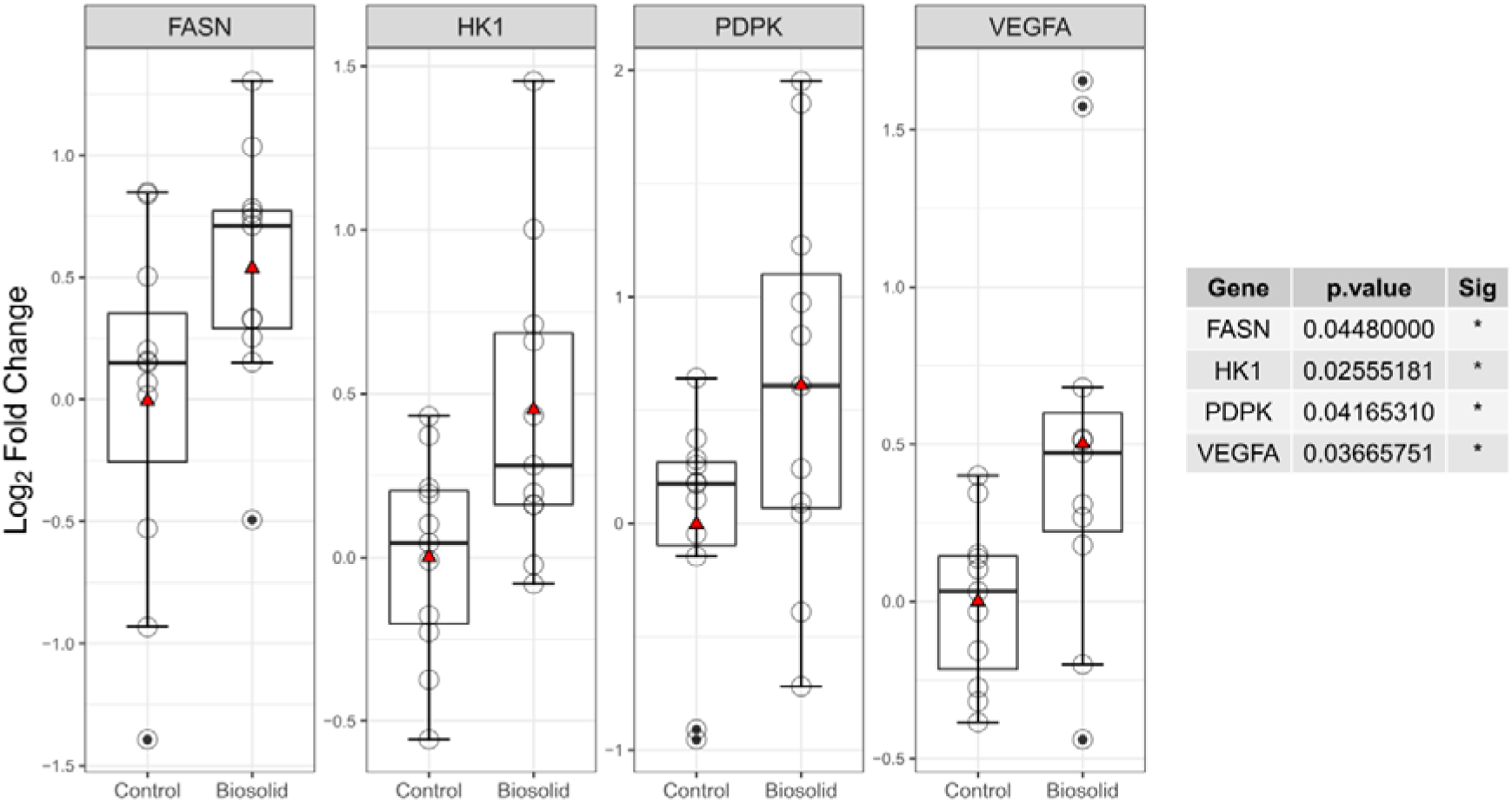
Log_2_ fold change of testicular gene expression between biosolid and control animals. Boxes represent 25” to 75^th^ percentile, horizontal bar indicates 50th percentile, whiskers indicate range excluding outliers, solid filled circles show outliers, open circles show individual data points, and red triangles show means.

### 2.5. Hypoxia Inducible Factor 1 Alpha analysis

Representative images from control and biosolid testes showing immunofluorescent staining for HIF1α and the nuclear marker DAPI are shown in Figure 4A, with the quantified overlap of HIF1α and DAPI signal (Manders’ overlap coefficient) shown in Figure 4B. In Leydig cells, a significantly (p = 0.0043) greater proportion of HIF1α was seen in the nucleus of biosolid (0.403 ± 0.106) compared to control (0.251 ± 0.097) animals (Figure 4B).

**Figure 4.**
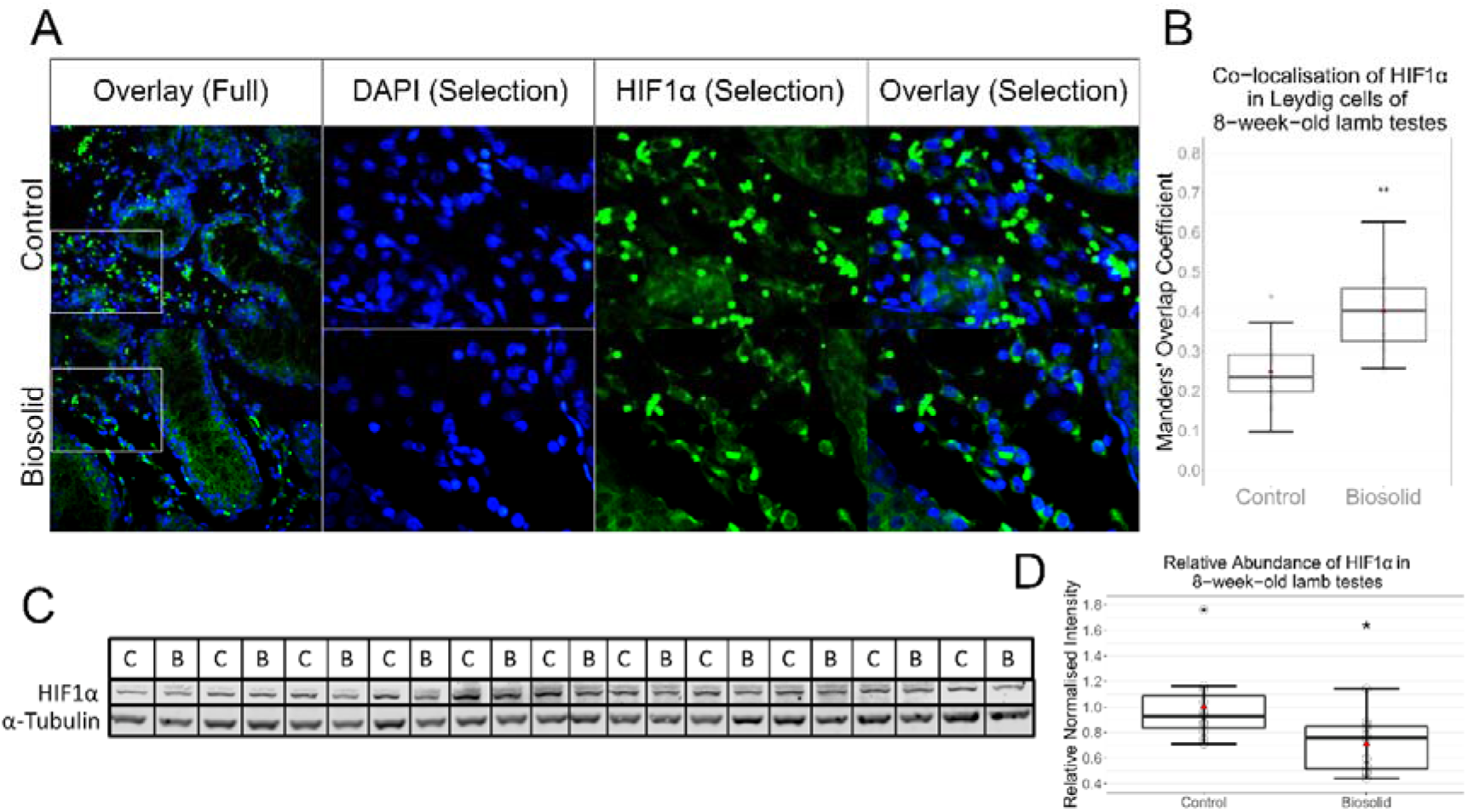
HIF1α in testes. Immunofluorescent staining of testes sections (A). Images to the right are of selections seen in images to the left. Blue staining is of DAPI and green of HIF1α. Co-localisation analysis (B) performed on regions outside of seminiferous tubules shows significantly (p = 0.0043) more nuclear localisation of HIF1α in biosolid animals than control. Manders’ Overlap Coefficients of the proportion of green signal overlapping with blue signal were determined by ImageJ. Western blots for HIF1α and a-Tubulin (C and D) show significantly (p = 0.0164) less HIF1α in biosolid animals than control. Boxes represent 25^th^ to 75^th^ percentile, horizontal bar indicates 50th percentile, whiskers indicate range excluding outliers, solid filled circles show outliers, open circles show individual data points, and red triangles show means.

Western blots for HIF1α and α-Tubulin are shown in Figure 4C with the mean relative abundance of HIF1α in the testes of control and biosolid animals shown in Figure 4D. There was significantly (p = 0.0164) less HIF1α detected in the testes of biosolid (0.710 ± 0.219) than control (1.00 ± 0.291) animals.

## 3. Discussion

The present study demonstrates the ability for gestational exposure to a low dose, extremely complex chemical mixtures to adversely affect prepubertal gonadal development. Similarities between the morphological and DGE patterns seen in biosolid exposed lambs and TDS patients suggest that the changes observed prior to puberty in this model may predispose animals to TDS and demonstrates the model’s utility to investigate the pathogenesis of TDS. Of specific note, biosolid exposure was associated with differential expression of genes related to apoptotic processes and mTOR signalling, and increased expression of a group of mTOR regulated genes which are all transcriptionally controlled by a common nuclear factor: HIF1α. The activation of HIF1α was also found to be altered in biosolid exposed animals visually, specifically where it was more localised to the nuclei of Leydig cells in biosolid exposed animals than in control. As HIF1α has previously been shown to inhibit STAR transcription and testosterone synthesis ^31^, this may be a mechanism through which gestational EC exposure may generate the altered testicular phenotype in biosolid exposed males, and may result in reduced fertility in affected animals.

There is increasing evidence of synergistic actions between different chemical components within low dose chemical mixtures producing adverse effects on the development of male gonads. Male offspring gestationally exposed to low doses of some simple chemical mixtures (containing ≤ 10 chemical components) have greater rates of feminised phenotypes, genital malformations, pathological testicular morphology, and impaired spermatogenesis ^11,32–35^. These studies exemplify the ability for even relatively simple chemical mixtures to elicit adverse effects in developing male gonads at low doses. In this study pregnant ewes were exposed to realistic levels and numbers of ECs as to be of human relevance. The present study showed anatomical differences between control and biosolid offspring, including lower bodyweights and pituitary weights in biosolid lambs. These results resemble anatomical differences seen in late gestation foetuses ^23^ and neonatal lambs ^24^, respectively, following similar exposure. The present study also showed fewer germ cells, with a concurrent increase in the occurrence of Sertoli-cell-only seminiferous tubules, in biosolid exposed lamb’s testes compared to controls, which phenotypically resembles mixed testicular atrophy – a hallmark of TDS ^2,36^. These results also resemble morphological differences seen in the testes of late gestation foetuses ^23^, neonatal lambs ^24^, and a subset of adult offspring ^17^, following similar exposure.

DGE analysis of the testicular transcriptome yielded many DEGs which may be involved in the pathogenesis of the observed phenotype. Noteworthy DEGs are a group involved in maintaining the blood-testes barrier (BTB) and cell polarity (*PRICKLE*, *CTNNA2*, and *DLG1*) ^37–39^ which had significantly lower expression in the testes of biosolid animals than in controls. BTB and cell polarity maintenance are crucial for germ cell stem cell survival and differentiation. Also noteworthy are a group of DEGs essential for spermatogenesis (*CIB1*, *GBF1*, *CFAP44*, and *PLK4*) ^40–43^, again with significantly lower expression in the testes of biosolid animals than in controls. This was also reflected in the GO term “Spermatid Development” being identified as significantly enriched within these DEGs. Comparison of the DEG data from this study with DEG data from human TDS patients allowed the identification of GO terms and KEGG pathways that were affected in both data sets, including two mTOR and two apoptotic entries. mTOR is known to be a crucial component of proper testicular development and spermatogenesis ^44^ and was previously indicated in transcriptomic analysis of testes from neonatal sheep gestationally exposed to BTP ^24^. Additionally, a range of chemicals have been shown to disrupt mTOR signalling pathways, causing testicular autophagy in pubescent rodents ^45–47^. A selection of genes were investigated which are known to have expression increased following mTOR activation ^48^. Of the four genes found to have higher expression in the testes of biosolid animals compared to controls, three (*VEGFA*, *HK1*, and *PDPK1*) are transcribed *via* Hypoxia Inducible Factor 1 Alpha (HIF1α) activation and contribute to the angiogenic and metabolic adaptive responses to hypoxia.

HIF1α is an important factor in embryonic development which is expressed from early embryonic stages, continuing in germ cells and other tissues into adulthood ^49^. In the classical scenario HIF1α responds to physiological oxygen status. Under normoxic conditions HIF1α is bound by the von Hippel-Lindau tumour suppressor protein (VHL) and degraded by ubiquitin-mediated proteolysis ^50^. Under hypoxic conditions the mechanisms for VHL association are suppressed and HIF1α will translocate to the nucleus to bind with HIF1β, forming the heterodimeric HIF-1 transcription factor, which binds to hypoxia-responsive elements (HREs) of target genes, increasing transcription ^51^. Separately to the classical scenario, HIF1α activation can occur independently of oxygen status by biochemical pathways (e.g. mTOR) ^52^ or small molecules and reactive oxygen species ^53,54^, with subsequent nuclear localisation, HIF-1 formation, and increased target gene transcription. Interestingly, contrary to other tissues, HIF1α can be stable under normoxic conditions in the testes. As HIF1α is expressed in germ cells throughout the life cycle, a role in germ cell development and maturation has been hypothesised ^49,55^, although precise details are yet to be discovered. However, it is Leydig cells which express the highest levels of HIF1α in the testes ^56^. The primary function of Leydig cells is testosterone production, which is tightly regulated in a closed-loop feedback mechanism by the hypothalamic-pituitary-testicular (HPT) axis. In steroid biosynthesis, the main rate-limiting step is the STAR mediated transport of cholesterol from the outer to inner mitochondrial membrane ^57^. HIF-1 has been shown to repress STAR transcription in Leydig cells by way of binding site blocking, and thus reduce testosterone synthesis ^31^. As greater nuclear localisation of HIF1α was observed in Leydig cells of biosolid animals compared to controls, further indicating HIF1α activation, this could provide a mechanistic reasoning to lower testosterone levels previously seen in neonatal biosolid lambs ^24^. However, hypoxia has also been reported to increase testosterone secretion by Leydig cells *in vitro* ^58^, which is potentially HIF1α mediated ^59^. It is also interesting that while nuclear localisation of HIF1α was greater within Leydig cells of biosolid animals, total testicular amounts of HIF1α were lower. However, this may be attributable to the changes in cellularity and the loss of germ cell HIF1α, and the magnitude of changes in localisation (160% of control) also outweighs the magnitude of changes in protein levels (71% of control).

Foetal development is an extremely complex and dynamic period, which increases vulnerability to xenobiotic induced toxicity. This is the first study to show similarity of DGE between low level EC exposure and human TDS patients. Additionally, this is the first study to link HIF1α activation to low level EC exposure. These findings add to the body of evidence to suggest that exposure to real-world levels of environmental chemical mixtures during pregnancy may be having an adverse effect on male offspring reproductive health, contributing to the decline in sperm quality and fecundity in humans.

## Abbreviations

BTP: Biosolid treated pasture
BPA: Bisphenol A
BTB: Blood-testes barrier
DGE: Differential gene expression
DEGs: Differentially expressed genes
EDCs: Endocrine disrupting chemicals
ECs: Environmental chemicals
FDR: False discovery rate
GO: Gene ontology
GnRH: Gonadotropin-releasing hormone
hCG: Human chorionic gonadotropin
HPT: Hypothalamic–pituitary–testicular
HIF1α: Hypoxia Inducible Factor 1 Alpha
HREs: Hypoxia-responsive elements
LH: Luteotropic hormone
NOAEL: No observable adverse effect level
PPCPs: Pharmaceuticals and personal care products
PBDEs: Polybrominated diphenyl ethers
PCBs: Polychlorinated biphenyls
PAHs: Polycyclic aromatic hydrocarbons
SCO: Sertoli-cell-only
TDS: Testicular dysgenesis syndrome
TDI: Tolerable daily intake
VHL: Von Hippel-Lindau tumour suppressor protein

## Acknowledgments

We are grateful to the staff at Cochno Farm and Research Centre for their technical assistance.

## 4. Online Methods

### 4.1. Ethics statement

All animals were maintained under normal husbandry conditions at the University of Glasgow Cochno Farm and Research Centre. The research programme was approved by the University of Glasgow School of Veterinary Medicine Research Ethics Committee. All procedures were conducted in accordance with the Home Office Animal (Scientific Procedures) Act (A(SP)A), 1986 regulations.

### 4.2. Experimental animals

EasyCare ewes were maintained on pastures fertilised with either biosolids, at conventional rates (4 tonnes/ha, twice per annum (April/September); biosolid), or with inorganic fertiliser at a rate which supplied equivalent levels of nitrogen (225 kg N/ha per annum; control) for one month prior to being mated and for the duration of pregnancy. Ewes were maintained indoor for the last two week of pregnancy and fed forage supplemented with concentrates as per normal husbandry practice, biosolids ewes received forage harvested from Biosolid treated pasture. Mating was by artificial insemination with semen from 4 rams which had only been maintained on control pasture. After parturition all ewes and lambs were maintained on control pastures, therefore all EC exposure was maternal i.e., through placental or lactational transfer. A subset of the male offspring from ewes exposed to conventionally fertilised pastures (n=11, control), and biosolid treated pastures (BTP) (n=11, biosolid), were weighed before euthanised by intravenous barbiturate overdose (140 mg/kg Dolethal, Vetroquinol, UK) for tissue collection.

### 4.3. Tissue collection

At post-mortem, gonads, thyroids, adrenals, and pituitaries were removed and weighed. Epididymides were removed from the testes and weighed. Testes weights were determined by subtraction of epididymides weights from gonadal weights. Two transverse slices were taken from the centre of the left testis, fixed overnight in 10% neutral buffered formalin (NBF, Thermo Scientific – 16499713), then transferred to 70% ethanol (VWR – 20821.330) prior to processing and embedding in paraffin wax for histology (Excelsior AS, Thermo Scientific). A 5mm transverse slice was taken from the centre of the right testis and frozen on dry ice prior to storage at −80°C until RNA extraction.

### 4.4. Immuno-histochemistry

Formalin-fixed paraffin embedded testicular tissue was sectioned (5μm) using a microtome (Leica Biosystems, model RM2125RT). Immuno-histochemistry was used to identify germ cells. One section per animal was mounted on a glass slide, dewaxed, and processed for antigen retrieval (autoclave for 21 minutes while immersed in citrate buffer 10mM, pH 6). Slides were washed in TBS and taken through peroxidase, avidin, and biotin blocking solutions (15 minutes each with TBS washes in between). Non-specific binding was blocked by incubation for 30mins with 20% goat serum in TBS before incubation with the primary antibody (rabbit anti-DDX4 polyclonal antibodies; Abcam – ab13840) diluted 1:1000 in antibody diluent (Agilent DAKO – S2022) overnight at 4°C. Sections were then washed in TBS + 1% Tween20 before being incubated with a biotinylated secondary antibody (goat anti-rabbit biotinylated polyclonal antibodies; Agilent DAKO – E0432) diluted 1:200 in antibody diluent for 30 minutes. Following incubation in secondary antibody, sections were treated with Vectrastain ABC-HRP system (Vector Laboratories – PK4000) for 60 minutes before being washed in TBS + 1% Tween20 and stained using DAB for 30 seconds. Slides were then washed in TBS, and counter-stained with haematoxylin and cover-slipped. Four representative images of the lobuli testis were captured (Leica DM4000B microscope with a Leica DC480 digital camera at 40x magnification using Leica Qwin software) from separate areas (top, bottom, left, and right) of each tissue section for each animal. Using ImageJ (version 1.53a), individual tubules which were entirely captured within images were manually selected (n = 2145 Control, 3673 Biosolid). DDX4 positive and negative cells were counted by automated macro (pre-validated on a subset of data – Supplementary Data 1). Mean germ cell to total cell populations, per tubule, were calculated relative to the mean control, as well as the proportion of seminiferous tubules without germ cells (Sertoli-cell-only; SCO). Separately, tubule selections were filtered for circularity using the equation *4π*(*Area*/*Perimeter*)^2 with a threshold of ≥0.9 (n = 587 Control, 1326 Biosolid) and minimum Feret’s diameters measured.

Fluorescent immuno-histochemistry was used to localise HIF1α. One section per animal was mounted on a glass slide, dewaxed, and microwaved at medium heat in 1 mM EDTA, pH 8.0, for 10 minutes. Sections were probed overnight at 4°C using mouse anti-HIF1α antibodies (Invitrogen – MAI-16504) diluted 1:250 in 5% BSA / TBS-T. Slides were washed with TBS and probed for 1 hour at room temperature using goat anti-mouse antibodies (Abcam – ab150113) diluted 1:1000 in 5% BSA / TBS-T. Sections were counter stained with DAPI (Abcam – ab104139) and cover-slipped. Four representative images of the lobuli testis were captured (Leica DM4000B microscope with a Leica DC480 digital camera at 160x magnification using Leica Qwin software) from separate areas (top, bottom, left, and right). Using ImageJ, areas out-with seminiferous tubules were manually selected and co-localisation analysis performed using the JACoP plugin ^60^, which provided Maders’ overlap coefficients for the proportion of HIF1α staining that overlapped DAPI staining.

### 4.5. RNA extraction, cDNA library preparation, sequencing, and data analysis

Transcriptome analysis was performed as previously described ^24^. Briefly, RNA was extracted, purified, reverse transcribed, and ligated to individual DNA barcodes for multiplexing. As the barcoding kit only contains twelve individual barcodes, samples were split into two groupings of eleven (mixed control and biosolids) for sequencing. Barcoded cDNA samples within a grouping were pooled and sequenced using a MinION Nanopore sequencer (Oxford Nanopore). Data were processed and filtered for quality prior to alignment to the reference transcriptome (constructed from NCBI’s Oar_v4.0 reference genome and annotation files) and counted. Differential gene expression (DGE) analysis was performed on gene counts by EdgeR. Differentially expressed genes (DEGs) were called using a p-value threshold of 0.05, log_2_ fold change threshold of <-1 or >1, and false discovery rate (FDR) threshold of 0.1. Gene ontology (GO) analysis was performed using DEG lists in DAVID (version 6.8).

### 4.6. BaseSpace Analysis

Illumina’s BaseSpace correlation Engine enables the comparison and correlation of DEG datasets through a combination of ranked-based enrichment statistics, meta-analyses, and biomedical ontologies ^61^. BaseSpace correlation Engine employs a rank-based, nonparametric analysis strategy driven by a Running Fisher’s test algorithm which performs the rank-based directional enrichment process. This enrichment process utilises a Fisher’s exact test to calculate four p-values: two p-values for the genes which are positively correlated between the datasets (genes that are either up or down-regulated in both datasets) and two for the negatively correlated genes (genes that are up-regulated in dataset 1 and down-regulated in dataset 2, or vice versa). The overall correlation p-value was calculated by converting the four p-values to −log_10_ p-values and subtracting the sum of the negative correlation p-values from the sum of the positive correlation p-values. A p-value threshold for significance of 0.0001 was used. To enable cross-platform and cross-species comparisons BaseSpace correlation engine software uses a database compiled of commonly used gene identifiers and reference identifiers along with ortholog information to standardise mapping across platforms and species.

Expression data for DEGs with p ≤ 0.05 were analysed for correlation to published data by Illumina’s BaseSpace software. Positively correlating DEGs were filtered for those common across each dataset per study. These gene lists were then combined and submitted to DAVID for pathway and GO analyses.

### 4.7. Effect of changes to cellularity

To assess the effect of changes in testicular cellular composition, DEGs with pathways or GO terms identified as enriched were tested for correlation using generalised linear models, on gene counts against the geometric mean of gene counts for germ cell specific biomarkers, and p values corrected for false discovery. Germ cell specific biomarkers used were CD9, CD14, THY1, NOTCH1, GFRA1, CDH1, and UCHL1.

### 4.8. Quantitative qPCR

RNA was extracted from approximately 30mg of frozen testes using RNeasy Mini Kit (Qiagen – 74104). Genomic DNA was degraded and cDNA synthesised using QuantiTect Reverse Transcription Kit (Qiagen – 205311). qPCR was performed using qPCR Brilliant II SYBR Master Mix (Agilent – 600828) on a Stratagene 3000. Primer details can be found in Supplementary Data 7. Raw fluorescent data were regressed by PCR Miner ^62^ to produce primer efficiencies and Ct values which were used in ΔΔCt analysis to produce log_2_ Fold Change values.

### 4.9. Western blots

Approximately 15 mg of frozen tissue samples were homogenisation using a 1 mL tapered PTFE tissue homogeniser into 19x volume of RIPA buffer (150 mM NaCl, 1% Triton X-100, 0.5% Sodium deoxycholate, 0.1% SDS, 50 mM Tris (pH 8.0)) containing protease inhibitors (Merck – 11697498001) and phosphatase inhibitors (Merck – 4906845001). Samples were centrifuged at 12,000 *xg* for 20 minutes at 4 °C. Following protein determination (Thermo Scientific – 23227), samples were made to a final concentration of 2 μg/μL using LDS Sample Buffer (Invitrogen – NP0007) and Reducing Agent (Invitrogen – NP0004) and heated to 95 °C for 10 minutes. 10 μL of reduced, denatured samples (20 μg protein) were loaded into wells of 4 to 12%, Bis-Tris, 1.0 mm acrylamide gels (Invitrogen – WG1403BOX) using 5 μL of protein reference standard (BioRad – 1610375EDU) in the first and last wells. Gels were run under constant voltage using MOPS SDS Running Buffer (Invitrogen – NP0001) with added antioxidant (Invitrogen – NP0005) and transferred to nitrocellulose membranes (Invitrogen – IB23001 NC) using an iBlot2 (Invitrogen – IB21001). Membranes were blocked using Intercept TBS Blocking Buffer (Licor – 927-60001), washed with TBS-T and TBS, and probed with overnight with primary antibody diluted 1:1000 in 5% BSA/TBS at 4 °C. The next morning membranes were washed with TBS-T and TBS before incubation in secondary antibody diluted 1:10,000 in 5% BSA/TBS at room temperature for one hour. Membranes were then washed in TBS-T, TBS, and finally MilliQ water before imaging on an Odyssey DLx Imager (Licor – 9142). Primary antibodies used were mouse-anti-HIF1α monoclonal (Invitrogen – MA1-16504) and rabbit-anti-α-tubulin polyclonal (Invitrogen – PA1-38814), with donkey anti-mouse (Invitrogen – SA5-10172) and donkey anti-rabbit (Invitrogen – SA5-10044) fluorophore conjugated secondary antibodies. Fluorescent intensities were quantified using Licor Image Studio Software (version 5.2.5). HIF1α signal intensities were normalised to α-Tubulin and expressed as relative to the average of control values.

### 4.10. Statistical analysis

All calculations and statistical analyses were performed in R (version 4.1.1) using base functionality. Unless otherwise stated, data were fitted to generalised linear models with gamma distribution and groups compared using one-way ANOVA. Plots were created using the R package ggplot2 (version 3.3.5). Data are presented as mean ± SD.

---

**Supplementary Data 1.**
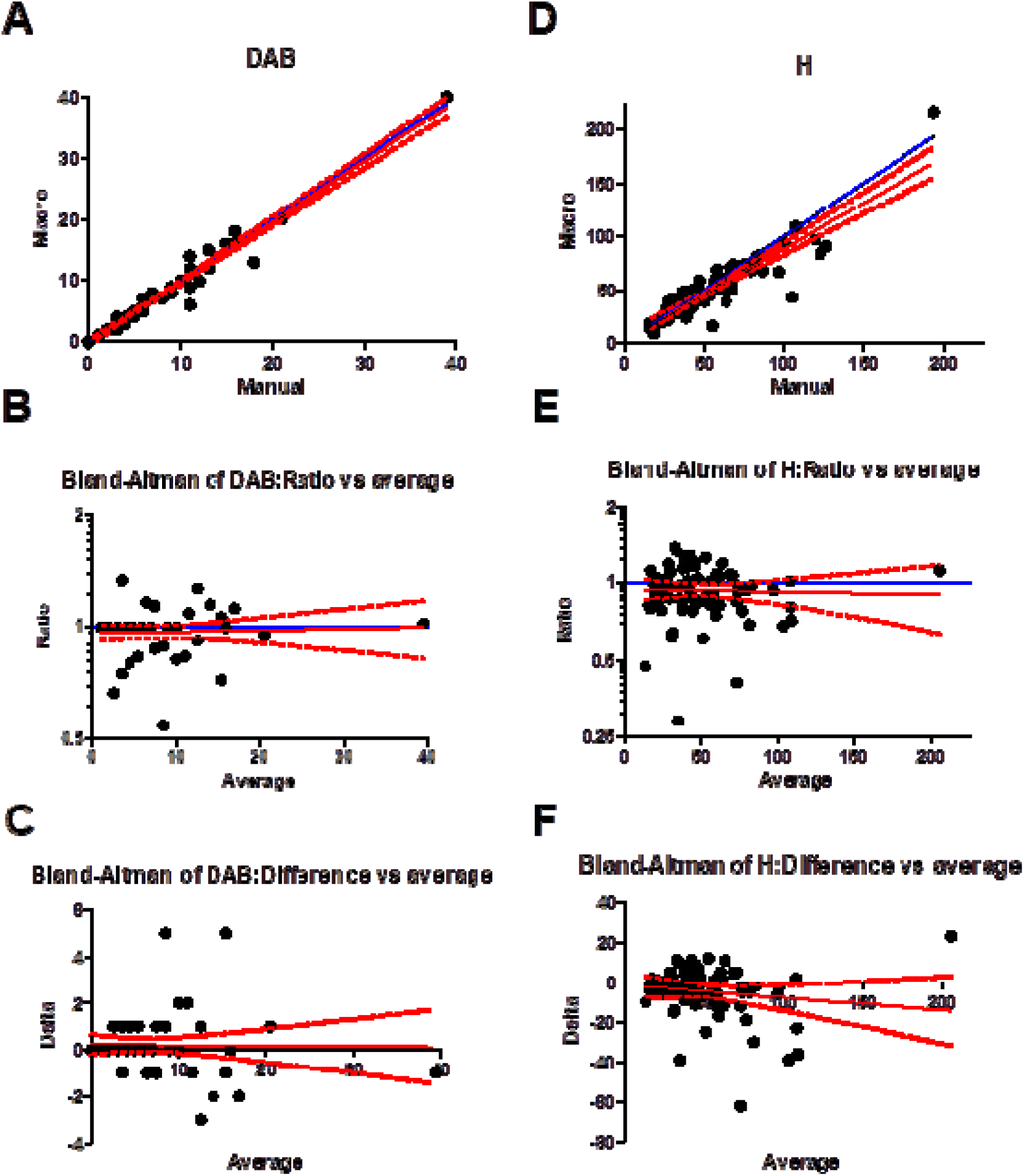
Comparative plots, and Ratio and Delta Bland-Altman plots for DAB (A, B, and C) and haematoxylin (D, E, and F) stained cells. Dots represent counts for individual seminiferous tubules. Blue lines represent theoretical complete agreement and red lines represent regressed lines of best fit (solid lines) and 95% confidence intervals (dotted lines).

**Supplementary Data 2.**
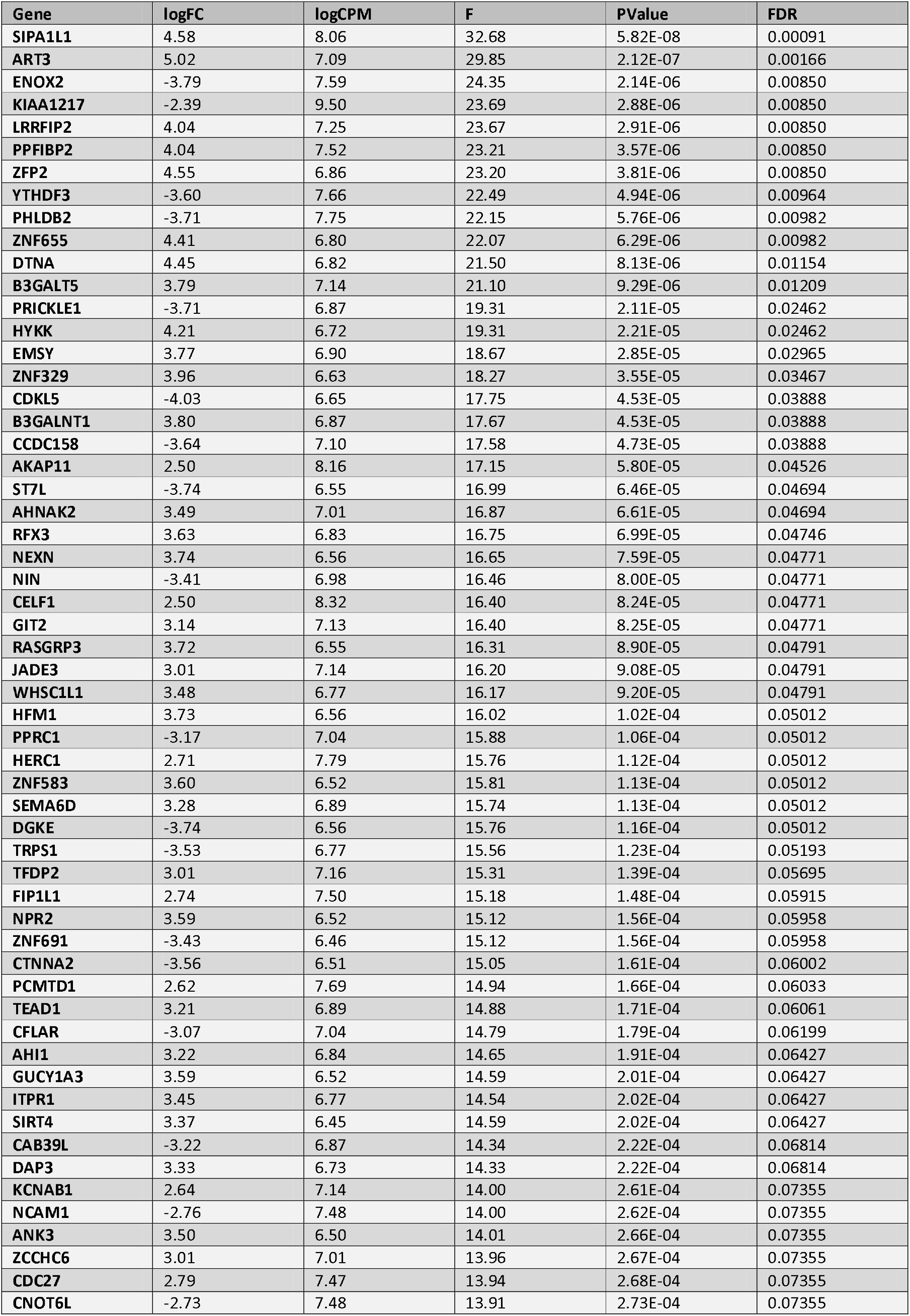
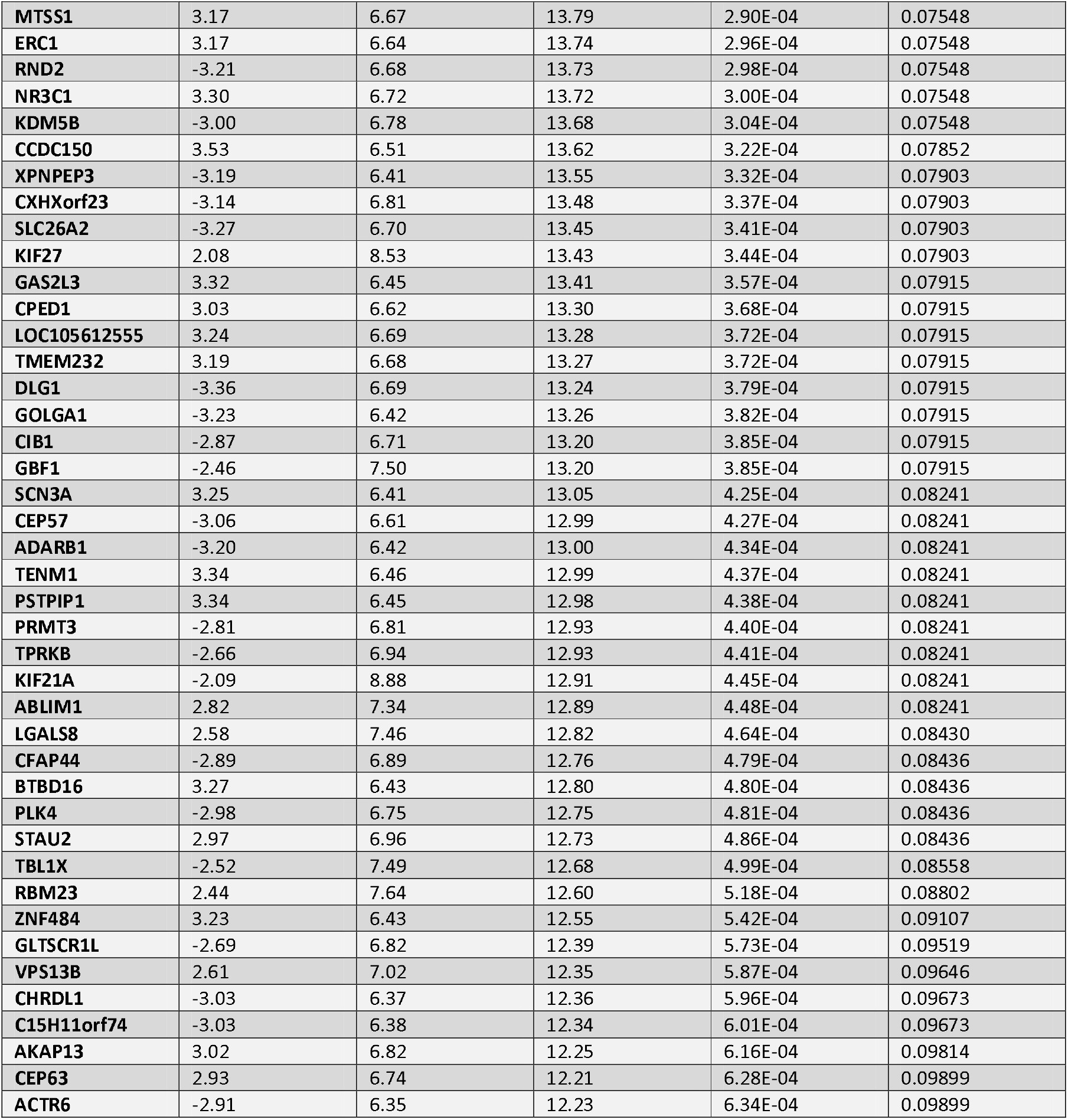
Differential expression data for the 50 genes with lowest p-values. LogFC = Log2 fold change between experimental conditions. LogCPM = Log2 counts per million. FDR = False discovery rate.

**Supplementary Data 3.** GO terms identified as enriched in the list of 99 DEGs with FDR < 0.1 by DAVID.

| Category | GO Term | Fold Enrichment | p-value |
| --- | --- | --- | --- |
| Cellular Component | Centrosome | 4.03 | 0.0156 |
| Molecular Function | Nucleic acid binding | 2.96 | 0.0160 |
| Biological Process | Spermatid development | 9.60 | 0.0379 |
| Cellular Component | Node of Ranvier | 47.01 | 0.0412 |
| Molecular Function | Guanylate cyclase activity | 45.21 | 0.0427 |
| Biological Process | Regulation of transcription, DNA-templated | 2.95 | 0.0498 |

Supplementary Data 4. Correlations between DEGs from biosolid vs control DGE analysis which were identified as having enriched GO terms, against the geometric mean of germ cell markers.

**Supplementary Data 5.** DEG data sets identified as having significant positive correlation with the biosolid DEG data.

| Name of Data Set | p-value | Number of Positively Correlating Genes | Number in Common | Number of Genes Combined |
| --- | --- | --- | --- | --- |
| Feig et al (2007) DOI: 10.1093/molehr/gal097 |  |  |  | 520 |
| Testicular biopsies of azoospermia patients with Sertoli-cell-only syndrome _vs_ full spermatogenesis | 3.57E-18 | 338 | 105 |  |
| Testicular biopsies of azoospermia patients with Sertoli-cell-only _vs_ spermatid stage arrest | 2.43E-16 | 274 |  |  |
| Testicular biopsies of azoospermia patients with Sertoli-cell-only _vs_ spermatocyte stage arrest | 2.21E-10 | 144 |  |  |
| Spiess et al (2012) DOI: 10.1093/humrep/dem292 |  |  |  |  |
| Testis from infertile man + Johnsen score 3.2 _vs_ full spermatogenesis | 3.44E-07 | 482 | 465 |  |
| Testis from infertile man + Johnsen score 2 _vs_ full spermatogenesis | 8.16E-06 | 542 |  |  |
| GSE6023 |  |  |  |  |
| SCOS patient testicular biopsy - Johnsen Score 10 _vs_ Score 2 | 2.80E-05 | 93 | 47 |  |
| SCOS patient testicular biopsy - Johnsen Score 9 vs Score 2 | 4.48E-07 | 98 |  |  |

Supplementary Data 6. Correlations between common DEGs from biosolid vs control DGE analysis and BaseSpace correlations which were identified as having enriched GO terms or pathways, against the geometric mean of germ cell markers.

**Supplementary Data 7.**
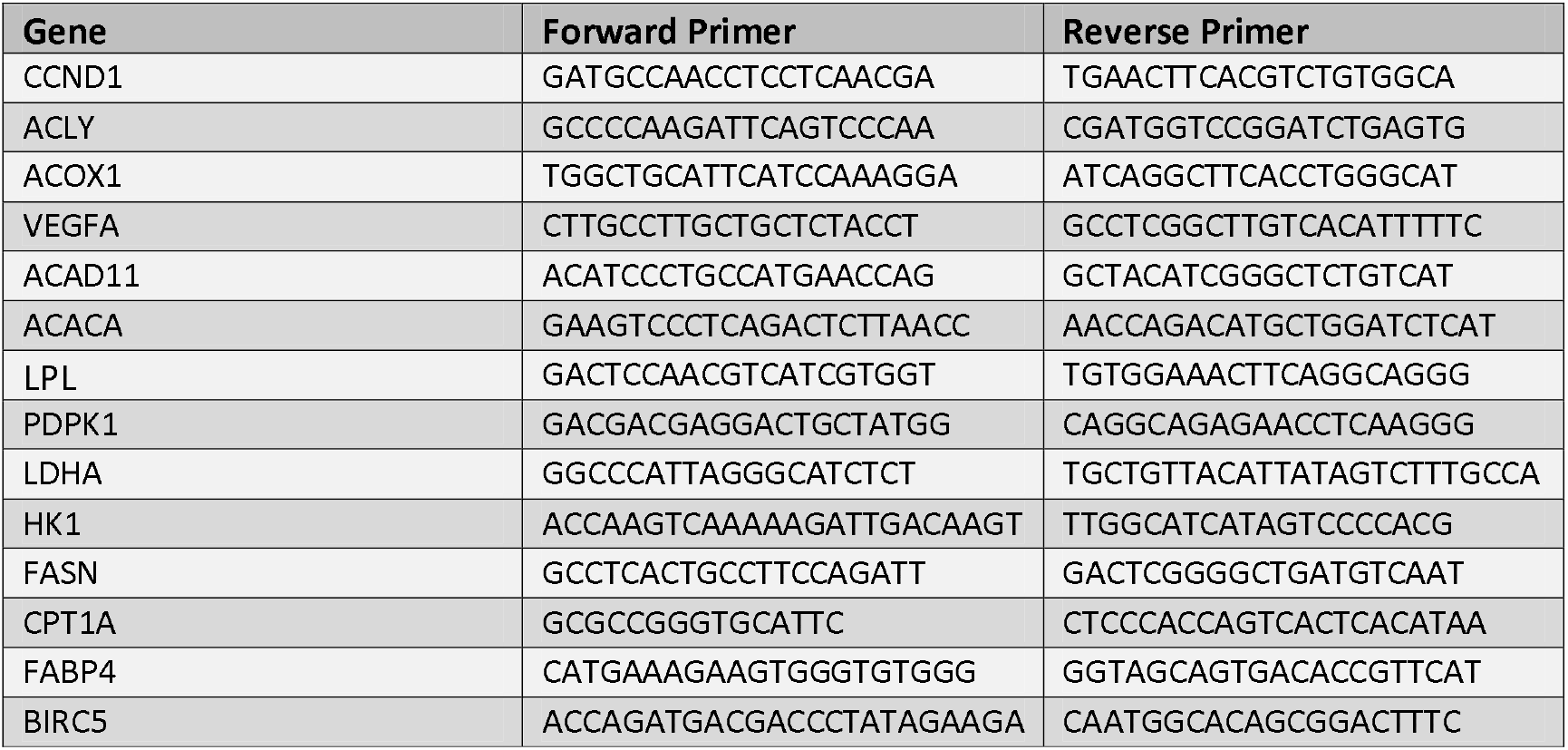
Primer details for qPCR.

## Notes

Funding: This work was supported by the National Institutes of Health [grant number R01 ES030374-01A1].

### Competing Interest Statement

The authors have declared no competing interest.

